# The Chicken Chorioallantoic Membrane as a Low-Cost, High-Throughput Model for Cancer Imaging

**DOI:** 10.1101/2023.06.21.545917

**Authors:** Lydia M. Smith, Hannah E. Greenwood, Will E. Tyrrell, Richard S. Edwards, Vittorio de Santis, Friedrich Baark, George Firth, Muhammet Tanc, Samantha Y.A. Terry, Anne Herrmann, Richard Southworth, Timothy H. Witney

## Abstract

**Purpose:** Mouse models are invaluable tools for radiotracer development and validation. They are, however, expensive, low throughput, and are constrained by animal welfare considerations. Here, we assessed the chicken chorioallantoic membrane (CAM) as an alternative to mice for preclinical cancer imaging studies.

**Methods:** Growth of NCI-H460 Fluc tumors on the CAM was optimized using a range of physical and chemical supports. Tumor-bearing eggs were imaged by dynamic ^18^F-2-fluoro-2-deoxy-D-glucose (^18^F-FDG) or (4S)-4-(3-^18^F-fluoropropyl)-L-glutamate (^18^F-FSPG) PET/CT following intravenous injection, with mice bearing subcutaneous NCI-H460 Fluc xenografts imaged with ^18^F-FDG for comparison. The dependence of the transporter system xc^-^ on *in ovo* ^18^F-FSPG tumor uptake was determined through treatment with imidazole ketone erastin. Additionally, ^18^F-FSPG PET/CT was used to monitor treatment response *in ovo* 24 h following external beam radiotherapy.

**Results:** NCI-H460 Fluc cells grown in Matrigel formed vascularized tumors of reproducible size without compromising embryo viability. By designing a simple method for cannulation it was possible to perform dynamic PET imaging *in ovo*, producing high tumor-to-background signal for both ^18^F-FDG and ^18^F-FSPG. ^18^F-FDG tumor uptake kinetics were similar *in ovo* and *in vivo*, with ^18^F-FSPG providing an early marker of both treatment response and target inhibition in CAM-grown tumors.

**Conclusions:** The CAM provides a low-cost alternative to tumor xenograft mouse models which may broaden access to PET and SPECT imaging. Rapid tumor growth and high-quality PET images that can be obtained with this model suggest its potential use for early radiotracer screening, pharmacological studies, and monitoring response to therapy.

## INTRODUCTION

The standard preclinical model for cancer research is the mouse. Multiple models have been developed using inbred mice to reflect the human disease, encompassing syngeneic, isogenic, spontaneous, and patient-derived tumors [1]. These models each have a unique set of advantages, with multiple models often used to answer a given research question. The wide-spread availability of xenograft mouse models has made it possible to study target expression, imaging agent specificity, metabolism, and pharmacokinetics at the organ and system level [2, 3] using preclinical PET scanners [4]. While powerful, such approaches have limitations. Tumor engraftment in mice can take many months and is complicated by variable take-rates, which increase costs. Additionally, animal housing units require extensive floor space and substantial investment for correct air handling, humidity, and temperature control. This, combined with the high price of immunodeficient animals and their associated husbandry costs, makes murine models unavailable to some [5]. Moreover, experiments must adhere to national animal licensing rules, which comes with further administrative responsibilities. Alternative models which resolve these ethical issues and reduce economic, time, and space requirements would therefore be highly advantageous.

Avian models may offer a viable alternative to mouse xenografts for preclinical cancer research. The fertilized chicken egg contains a highly vascularized membrane that surrounds the embryo, named the chorioallantoic membrane (CAM), which provides an ideal environment for tumor growth [6, 7]. The CAM is formed when the chick allantois and chorion fuse on embryonic day 6-7 (E6-7) [8]. The CAM performs gas exchange and provides nutrients for the chick’s growth by drawing calcium from the porous shell [9]. In this immunodeficient, vascularized, and oxygen-rich environment, human tumors require just 3-7 days to establish [10]. Consequently, this model has been used to study tumor growth [6, 11], metastasis [12, 13], and angiogenesis [14, 15]. Moreover, the CAM has sufficient optical transparency for intravital microscopy, allowing the observation of cell migration within the vasculature [16]. Other uses include the assessment of drug delivery [17] and screening of therapeutic agents [18]. Importantly, the chick CAM aheres to the principles of the 3Rs (Replacement, Reduction and Refinement) [19] as it is not recognized as a protected species until E14 under European law (directive 2010/63/EU).

To-date, relatively few studies have used the chick CAM to evaluate novel radiopharmacuticles. Previously, PET/CT to assess glucose uptake and the proliferation of tumors grown on the chick CAM using ^18^F-2-fluoro-2-deoxy-D-glucose (^18^F-FDG) and ^18^F-Fluorothymidine (^18^F-FLT), respectively [14]. From the resulting images, it was possible to delineate radiotracer uptake in both the tumor and chick [14]. Other studies have investigated bone metabolism with ^18^F-fluoride [20] and the tryptophan metabolic pathway with 7-^18^F-fluorotryptophan in the chick CAM, where uptake mechanisms and levels of dehalogenation were found to be comparable to the mouse [21]. More recently, ^18^F-siPSMA-14 PET/MRI was used to image PSMA-positive and -negative tumors both *in ovo* and *in vivo* [22], with ^68^Ga-Pentixafor PET/MRI used to evaluate colorectal cancer uptake and blocking with a CXCR4 antagonist in addition to ^18^F-FDG, which was used as a viability marker [23]. Current unresolved limitations with this model relate to variable tumor growth rates, difficulties cannulating the CAM vessels for dynamic imaging, and the need to cool the egg to immobilize the embryo, which negatively impacts radiotracer delivery, internalization, and metabolism [22].

To be widely-adopted as a model for radiopharmaceutical research, a straight-forward protocol for the use of the chick CAM must be established. Here, we present a simple method for vessel cannulation and the use of liquid narcotics for chick immobilization. With this optimized protocol and through direct comparison studies, we show that the chick CAM is a suitable intermediate that may precede more complex experimental models, which may reduce the reliance on subcutaneous tumor-bearing mice. As well as investigating the optimal conditions for *in ovo* non-small cell lung cancer (NSCLC) growth, we assessed *in ovo* and *in vivo* tumor imaging and pharmacokinetics using ^18^F-FDG and the system xc^-^ substrate (4S)-4-(3-^18^F-fluoropropyl)-L-glutamate (^18^F-FSPG). Finally, we asked whether this model could be used for other applications, such as target engagement studies using the system xc^-^ inhibitor imidazole ketone erastin (IKE), and to determine early response to external beam radiation.

## METHODS

### Radiochemistry

Clinical-grade ^18^F-FDG and ^18^F-fluoride was acquired from King’s College London & Guys and St Thomas’ PET Centre. ^18^F-FSPG radiosynthesis (GE FASTlab) and quality control was performed according to previously-published methodology [24].

### Cell culture

NCI-H460 Fluc (PerkinElmer) were cultured in RPMI 1460, supplemented with 10% fetal bovine serum, 25 mM L-glutamine, 100 U.mL^−1^ penicillin and 100 µg.mL^−1^ streptomycin (ThermoFisher Scientific). All cells were maintained in a humified atmosphere at 37 °C and 5 % CO2.

### Tumor engraftment

Prior to chick CAM tumor cell inoculation, fertilized Dekalb white eggs (Henry Stewart & Co. Ltd) were stored at 12 °C. To initiate embryo growth, E0 eggs were moved into an Ovaeasy 190 advance EX series II incubator (Brinsea), running at 37.6 °C and 50% humidity and set to tilt every 30 min. On E3, eggs were removed from the incubator and windowed following previously-defined protocols [25] then placed in a second incubator set to 37.6 °C and 50% humidity with no tilt setting. On E7, eggs were removed and inoculated with 3 × 10^6^ NCI-H460 Fluc cells with a range of chemical and physical supports (see below). The eggs were then placed back in the incubator and kept until E14. In some instances, eggs formed two tumors of a similar size. These eggs were used for target engagement studies as the second tumour could be used as an intrinsic control.

### Optimization of *in ovo* tumor growth

To determine the best method for tumor innoculation, five sets of 3 × 10^6^ NCI-H460 Fluc cells were harvested on E7 (500 × *g* for 3 min at 4 °C), supernatants discarded, and cell pellets kept on ice. Tumors were grown on the CAM according to the following conditions: Group 1 (*n = 10*), cell pellet with 5 µL trypsin applied onto the CAM prior to injection; Group 2 (*n = 14*), cell pellet mixed in 20 µL Matrigel (Corning); Group 3 (*n = 13*), cell pellet in 20 µL growth factor reduced (GFR) Matrigel (Corning); Group 4 (*n = 11*), cell pellet in 20 µL complete medium; Group 5 (*n = 13*), cell pellet injected in the middle of a 12 mm diameter sterile plastic ring laid on top of the CAM; and Group 6 (*n = 11*), cell pellet injected into the albumin below the surface of the CAM. Embryo survival to E14 and the tumor take-rate of each group was recorded. Tumor sizes were measured by excizing and recording their wet weight on E14.

### Cannulation optimization

Initially, a 30g insulin syringe was used for direct injection into the CAM vessels (**Fig. 1a**), without success. Next, a 30g insulin syringe was cut to form a short needle and cannula tubing was pulled over the end. The needle and tubing was held with needle holders (World Precision Instruments) and an injection by hand was performed (**Fig. 1b**). Again, there was limited success with this method. Then a micromanipulator (Prior, UK) mounted with a curved spatula was used to provide a solid support for the vessel outside of the eggshell, thereby overcoming the restrictions imposed by cannulating *in situ*. The vessel was accessed by cutting the CAM either side of the vessel and hooking the spatula underneath. The micromanipulators were then raised to tension the vessel. An injection by hand was performed using the cut 30g needle, tubing and needle holders (**Fig. 1c**). To further improve this technique, glass needles were made using a needle puller, model PP-830 (Narishige), where 1.17 mm borosilicate glass capillary tubes were heated to 60 °C for 35 s before being pulled into a sharp point. Peristaltic pump tubing was placed over the end of the capillary tube to form a cannula. Using the micromanipulator set up described previously, an injection into the tensioned vessel was performed by hand and the needle tied in place with a suture (**Fig. 1d**). Lastly, cannula tubing was inserted through the glass needle and glued in-place. This cannula was used to inject at a branch point of the vessels by hand and the needle secured in place with vetbond glue (**Fig. 1e**) and was used for all imaging experiments (**Fig. 1f & g**).

**FIGURE 1.**
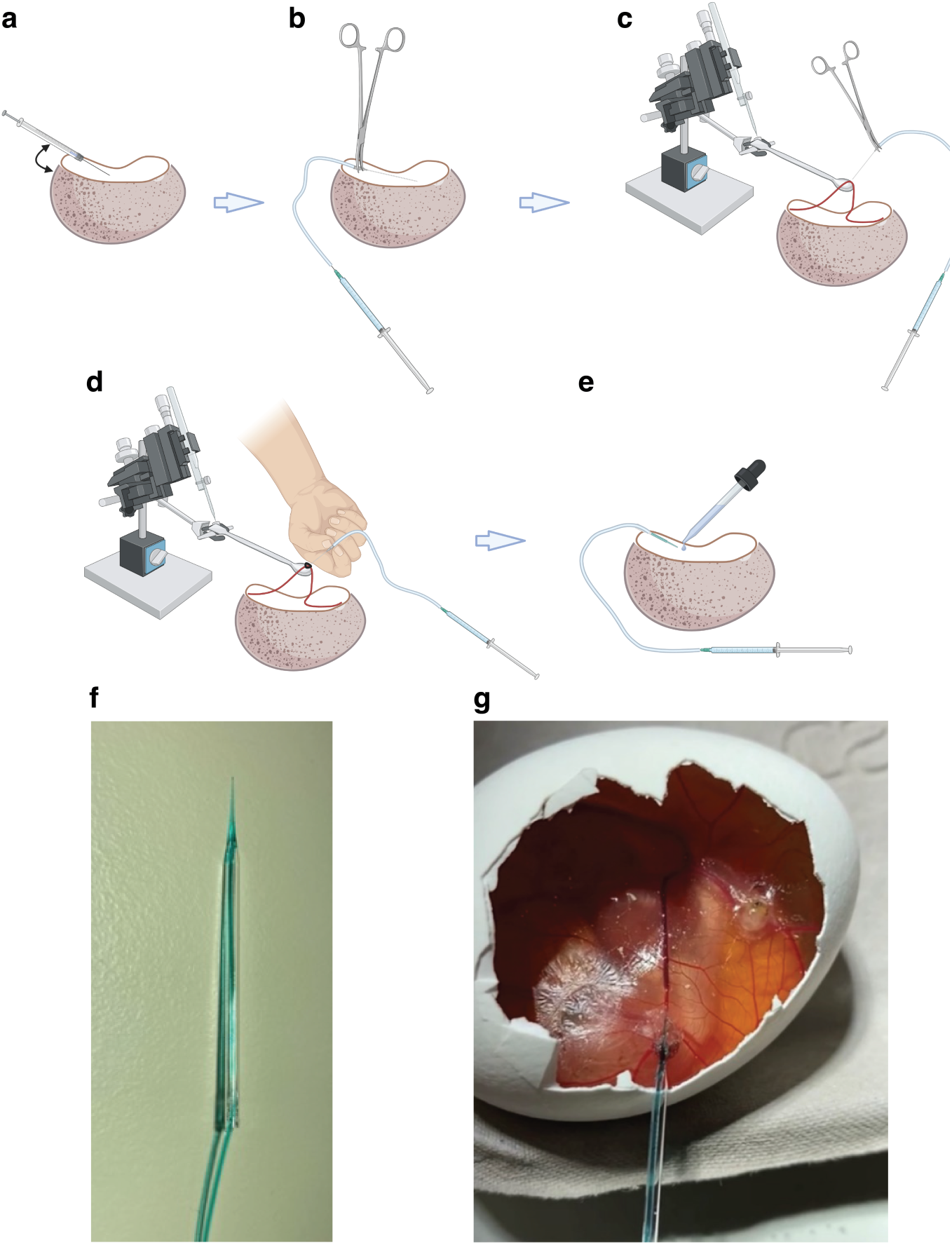
Evolution of Chick CAM cannulation methodology. Various methods were trialed during cannulation optimization: a. Direct injection with 30g insulin syringe by hand. b. Cut 30g needle with needle holders. c. Micromanipulators fixed with a hooked spatula attachment used to pull the vessel over and provide tension. d. Glass needle and peristaltic pump tubing tied in with suture. e. Glass needles injected by hand at branch points and secured in place using vetbond glue (optimized technique). f. Photo of the final glass needle cannula used for all subsequent experiments. g. Successful cannulation of a CAM vessel using (f).

### Intravenous injection

Multiple experimental setups were investigated for CAM vein cannulation (illustrated in **Fig. 1**). In the final optimized procedure, glass needles were made using a needle puller (model PP-830, Narishige) where 1.2 mm borosilicate glass capillary tubes were heated to 60 °C for 35 s before being pulled into a sharp point. Cannula tubing was inserted through the glass needle and fitted to the end of the needle to reduce dead volume. The tubing was glued in place to make a cannula. Glass needles were used to inject at a branch point of the vessels by hand and secured in place with vetbond glue. Successful cannulation was confirmed by injection of ∼20 µL of fast green dye (Sigma).

### Immunofluorescence

On E14 NCI-H460 Fluc tumor-bearing eggs were cannulated and injected with a bolus (∼200 µL) of Hoescht 33342 (1:200 in PBS, Invitrogen) and 0.05 mg/mL lens culinaris agglutinin (1:20 in PBS, 2BScientific, DL-1048). Following 10 min, the tumors were excised and coated in optimal cutting temperature compound (OCT, ThermoFisher Scientific) and placed in isopentane cooled with liquid nitrogen until the OCT had set. Tumors were then transferred to liquid nitrogen for a further minute prior to storage at −80 °C. 20 µm tumors sections were cut at −18 °C using a cryostat (SLEE) and mounted onto a microscope slide. The tissue was fixed by submerging the slides in 3% formaldehyde for 30 min, followed by rehydration with PBS. Tissue was permeabilized with 1% BSA, 0.1% Titron in PBS applied for 30 min. Next, blocking solution was added (5% goat serum and 5% BSA in PBS) for 20 min. Cytokeratin 18 antibody (abcam) was prepared (1:200 in 5% BSA) and 30 µL was applied to each slice of tissue before slides were left in a dark humidified box overnight at 4 °C. The following day, slides were washed three times with phosphate-buffered saline with Tween-20 (PBST), with 30 µL GFP-labelled secondary antibody (abcam, 1:200 in 5% BSA) then added to each tissue slice. Slides were left to incubate in the dark for 2 h. Finally, slides were washed with PBST before a drop of EverbriteT hard set mounting medium without DAPI (Cambridge Bioscience) was applied with a coverslip to set the slides. Slides were imaged at the KCL Nikon Imaging Center using a spinning disc confocal microscope (Nikon).

### Bioluminescence imaging

*In ovo* tumors were imaged by bioluminescence imaging (BLI) using an IVIS Spectrum *in vivo* imaging system (PerkinElmer). Images were acquired seven days post cell inoculation to confirm successful tumor growth. 1 g of firefly luciferin (Promega) was dissolved in 66.6 mL of sterile PBS to create a 15 mg/mL solution. The luciferin was sterile filtered through using a 0.2 µm filter and aliquoted into 5 mL vials for freezing. An *i.v.* injection of 200 µL of 15 mg/mL firefly luciferin was performed before the egg was transferred to the IVIS Spectrum camera and maintained at 37 °C. Images were acquired until the luminescent signal plateaued ∼20 min p.i. of luciferin, ensuring maximum tumor signal was reached (exposure time 1-60 s, binning 2-8, FOV 23 cm, f/stop 1, no filter). For signal quantification, images were analyzed using Living Image software (PerkinElmer). A region of interest was drawn around the tumor to measure total photon flux (photon/sec).

### PET imaging of *in ovo* tumors

Eggs bearing NCI-H460 Fluc tumors were grown in Matrigel, as described above. On E14, a CAM vein was cannulated and 90 µL of a 1 mg/mL solution of the anesthetic medetomidine was pipetted on to the surface of the CAM. Eggs were left for 15 min at room temperature before receiving a bolus injection of ∼3 MB ^18^F-FDG (*n* = 7) or ^18^F-FSPG (*n* = 10) on the imaging bed (<150 µL) and washed through with 50 µL PBS. 60 min dynamic or 20 min static PET scans 40-60 min post injection (p.i.) were acquired using a Mediso NanoScan PET/CT system (1-5 coincidence mode, 3D reconstruction, attenuation-corrected, scatter-corrected). CT images were obtained for attenuation correction (180 projections, semi-circular acquisition, 50 kVp, 300 ms exposure time). The eggs were kept at 37 °C throughout the scan. Dynamic PET data were reconstructed into 19 bins of 4 × 15 seconds, 4 × 60 seconds, and 11 × 300 seconds (Tera-Tomo 3D; 4 iterations, 6 subjects, 400-600 keV, 0.3 mm^3^ voxel size). VivoQuant software (v2.5, Invicro Ltd.) was used to analyze the reconstructed images. Regions of interest (ROIs) were drawn manually using 40-60 min summed PET images. Finally, time verses radioactivity curves (TACs) were generated, and, area under time verses radioactivity curves (AUC) were calculated. For inhibition studies, eggs bearing NCI-H460 Fluc tumors received an intratumoral injection of IKE (2.5 mg/kg, in 5% DMSO, 95% PBS) 60 min prior to PET imaging.

### Biodistribution

*Ex vivo* biodistribution studies were performed on eggs bearing NCI-H460 Fluc tumors (*n* = 6) one h p.i. following an *i.v.* injection of ^18^F-FSPG (0.5 MBq, 200µL). Chicks were culled by direct injection of 50 µL of pentobarbital (200 mg/mL) and tissues of interest were collected. All tissues were washed in phosphate buffered saline and weighed. A gamma counter was used to count the tissue (1282 compugamma, LKB; window set to channels 175-220 for the energy profiles). *Ex vivo* biodistribution data were presented as %ID/g.

### Tumor irradiation

At E13 eggs were randomized into CT only (delivering 0.12 Gy, *n* = 6) and irradiated groups (Precision X-Ray, Inc. SmART+ small animal irradiator, *n* = 6). Eggs were scanned by cone beam CT with a 2 mm aluminium filter using the mouse soft tissue, high dose parameters at 40 kVp and 12 mA, producing 0.1 mm voxels for Monte Carlo treatment planning using the SmART-ATP software (v2.0.20201216). The ROIs were hand-drawn and interpolated to create a 3D volume of interest. The isocentre and collimation were set to minimize irradiation of the chick embryo using parallel-opposed pair of beams at 0 and 180 degrees, which were optimized using Monte Carlo simulations (100 million photon histories) which modelled the mean radiation dose delivered to the isocentre (D mean) and normal tissues. Radiation was delivered as a single 12 Gy fraction to the tumor using a circular collimator and a 0.3 mm Cu filter. After irradiation and/or CT imaging, eggs were placed back into the incubator for 24 h prior to PET imaging.

### Studies in mice

All experiments in mice were performed in accordance with the United Kingdom Home Office Animal (scientific procedures) Act 1986.

### *In vivo* NCI-H460 Fluc tumor growth & BLI imaging

A suspension of 100 µL PBS containing a total of 3 × 10^6^ NCI-H460 Fluc cancer cells was injected subcutaneously into female Balb/c nu/nu mice aged 6 to 9 weeks (Charles River Laboratories, *n* = 20). Tumor dimensions were measured using calipers and the volume calculated using the following equation: volume = [(*π*/6) × *h* × *w* × *l*], where *h, w*, and *l* represent, height, width, and length, respectively. The mice bearing subcutaneous NCI-H460 Fluc tumors were also imaged by BLI using an IVIS Spectrum *in vivo* imaging system (PerkinElmer) to confirm successful implantation. Mice were subsequently monitored and imaged twice a week for three weeks or until experimental end point. Luciferin was prepared as stated above. Prior to imaging, mice were anesthetized with isoflurane (2 % in O2) and injected with an i.p. injection of 200 µL of firefly luciferin. Mice were then transferred to the IVIS Spectrum camera and maintained at 37 °C. Images were acquired until the luminescent signal plateaued ∼20 min p.i., ensuring maximum tumor signal was reached (exposure time 1-60 s, binning 2-8, FOV 23 cm, f/stop 1, no filter). For signal quantification, images were analyzed using Living Image software (PerkinElmer). A region of interest was drawn around the tumor and total photon flux was measured (photon/sec). Mice were selected for PET/CT imaging once the bioluminescent signal reached ∼2.9 × 10^9^ photons/s/cm^3^ or the tumors reached approximately 100 mm^3^.

### ^18^F-FDG PET imaging of mice bearing NCI-H460 Fluc tumors

Dynamic 60 min ^18^F-FDG PET scans were acquired on a Mediso NanoScan PET/CT system (1-5 coincidence mode; 3D reconstruction; CT attenuation corrected; scatter corrected) following a bolus *i.v.* injection of approximately 3 MBq of ^18^F-FDG (<200 µL) into mice bearing subcutaneous NCI-H460 Fluc tumor xenografts (*n* = 9). Mice were kept at 37 °C throughout the scan. CT images were obtained for attenuation correction (180 projections; semi-circular acquisition; 50 kVp; 300 ms exposure time). The acquired PET data was reconstructed into 19 bins of 4 × 15 seconds, 4 × 60 seconds, and 11 × 300 seconds (Tera-Tomo 3D reconstructed algorithm; 4 iterations; 6 subjects; 400-600 keV; 0.3 mm^3^ voxel size). VivoQuant software (v2.5, Invicro Ltd.) was used to analyze the reconstructed images. ROIs were drawn manually using the CT image. Images were processed as described above. At the end of the scan, tumors were excised and snap-frozen in liquid nitrogen for *ex vivo* analysis.

### GSH assay

*In ovo* tumors were collected and lysed in the buffers for the GSH/GSSG-Glo assay kit (Promega) according to the manufacturer’s instructions and normalized for protein concentration (Pierce BCA protein assay kit, ThermoFisher Scientific).

### Western blotting

Western blot analysis was carried out on cell, *in ovo* and *in vivo* tumor lysates using an established experimental method described in [26]. The protein concentration of samples was determined using the Pierce BCA protein assay kit. For these experiments all antibodies were purchased from Cell Signaling

Technology and were anti human antibodies raised in rabbit at 1:100 dilution. A HRP linked anti rabbit secondary antibody was used to visualize with an iBright imaging system (ThermoFisher Scientific).

### H&E staining

Tumors excised at E14 were submerged in 70% ethanol overnight, followed by 95% ethanol for a further 2 h. Next, tumor were placed in 100% ethanol for 2 h, followed by xylene for 90 mins, prior to being paraffin embedded. Embedded tissue was stored at 4 °C until being processed by UCL IQPath for histologic analysis.

### Statistics

GraphPad Prism (v.8.0) was used to perform statistical analysis on data. All data were expressed as the mean ± standard deviation (SD). Statistical significance was determined using either unpaired or paired two-tailed Student’s t-test for data that fit the category of parametric analysis or Mann-Witney U test for data which required a nonparametric analysis. For analysis across multiple samples, 1-way analysis of variance (ANOVA) followed by multiple comparison correction (Tukey) was performed. Groups were considered significantly different from each other if *p* < 0.05.

## RESULTS

### Matrigel is the optimal matrix for *in ovo* tumor growth

Embryo viability was assessed on E14 following NCI-H460 Fluc tumor cell inoculation using a range of physical and chemical supports (**Fig. 2a)**. When RPMI was used alone, survival rates were <50 %. Embryo survival increased to 71, 77 and 88% with the use of Matrigel, GFR Matrigel or CAM pretreatment with trypsin, respectively (*n* = 11-14). The use of a ring or albumin as a tumor support had a negative impact on embryo survival, with only 16 % and 36 % surviving to E14, respectively. RPMI, Matrigel, GFR Matrigel, and the trypsin group achieved a tumor take-rate of ∼80%, while only 50% was achieved with the ring support. Direct injection of cells into the albumin did not result in tumor formation (**Fig. 2b**). GFR Matrigel mixture gave the largest, but most variable tumors on average (0.11 ± 0.06 g). While non-GFR supplemented Matrigel produced smaller tumors, they were more consistent in size (0.09 ± 0.02 g; **Fig. 2c**) and location of growth (**Supplementary Fig. 1**). Matrigel was therefore selected as the growth matrix for subsequent PET imaging experiments. The viability CAM-grown NCI-H460 Fluc tumors was visualized by BLI (**Fig. 2d**) and the absence of apoptosis was confirmed by measurements of cleaved caspase 3 (**Supplementary Fig. 2**). Hematoxylin and eosin (H&E) and immunofluorescent staining revealed a heterogenous, but well vascularized and perfused tumor (**Fig. 2e** & **f**).

**FIGURE 2.**
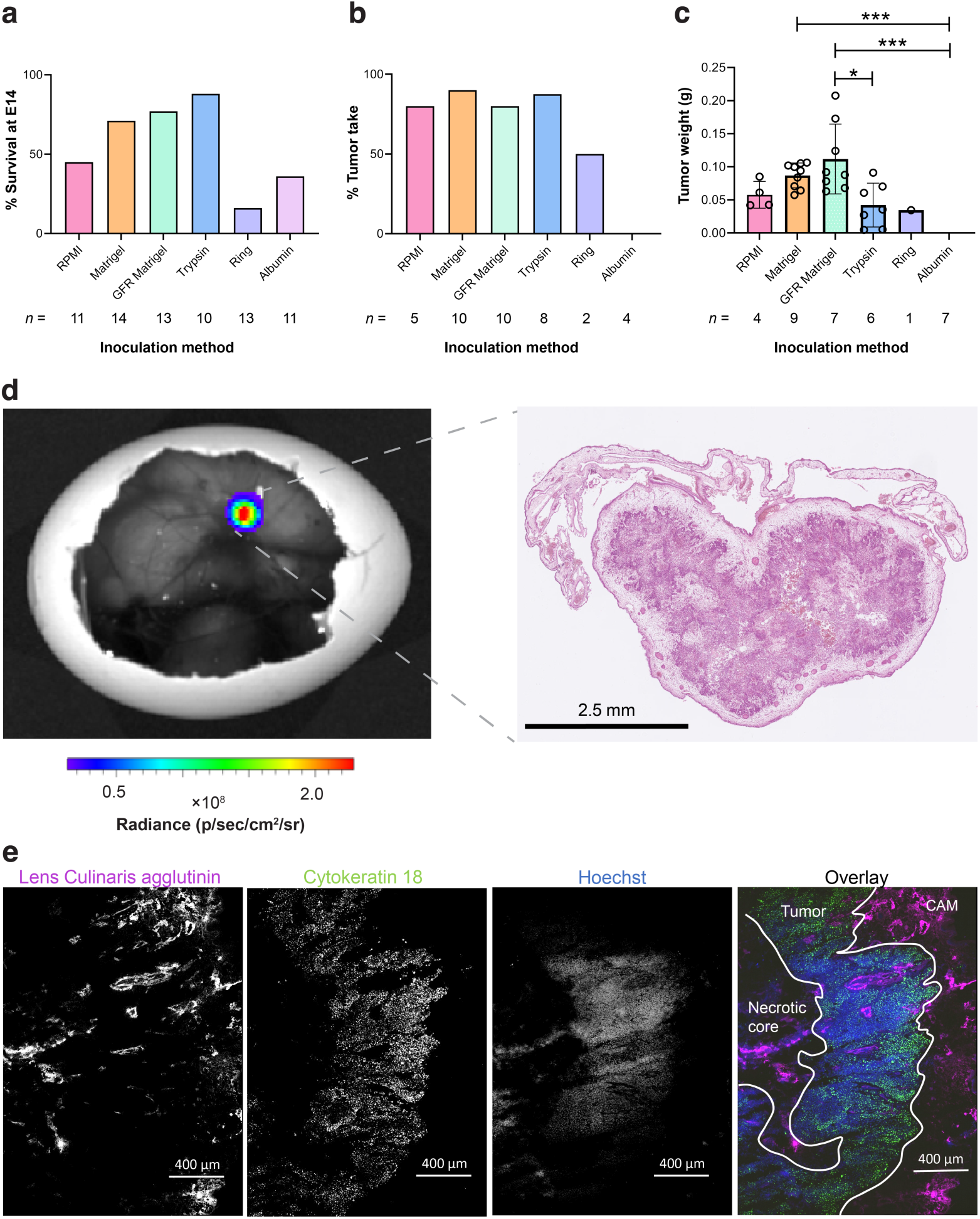
Optimization and characterization of *in ovo* tumor growth. a. Relative chick survival on E14 using a range of chmical and physical supports. b. Percentage tumor take-rate of surviving embryos from each inoculation group. c. Tumor weight at E14. d. Representative BLI image of an *in ovo* NCI-H460 Fluc tumor. e. Accompanying H&E section. f. Separate and overlay images showing vasculature (lens culinaris agglutinin, 649 nm), tumor cell cytoskeleton (cytokeratin 18, 488 nm) and perfusion with nuclear stain Hoechst 33342 (350 nm). Error bars show standard deviation. *, *p* < 0.05; ***, *p* < 0.001.

### ^18^F-FDG tumor pharmacokinetics are comparible *in ovo* and *in vivo*

To determine whether CAM-grown tumors were a viable alternative to mouse xenografts for molecular imaging applications, we performed dynamic ^18^F-FDG PET/CT imaging of NCI-H460 Fluc tumors *in ovo* (**Fig. 3a** and **Supplementary Fig. 3a**). Unlike in mice (**Fig. 3b**), ^18^F-FDG was homogenously distributed throughout the embryo, with no clear pattern of radiotracer clearance. *In ovo*, ^18^F-FDG uptake was highest in the tumor, which was characterized by rapid initial delivery, reaching 7.0 ± 0.8 %ID/g at 5 min, followed by a slower rate of uptake (10.0 ± 2.1 %ID/g at 60 min, **Fig. 3c**). We selected the yolk sac as the background ROI which was 0.7 ± 0.6 %ID/g at 60 min (**Supplementary Fig. 3b**), giving a tumor-to-background ratio of ∼15.

**FIGURE 3.**
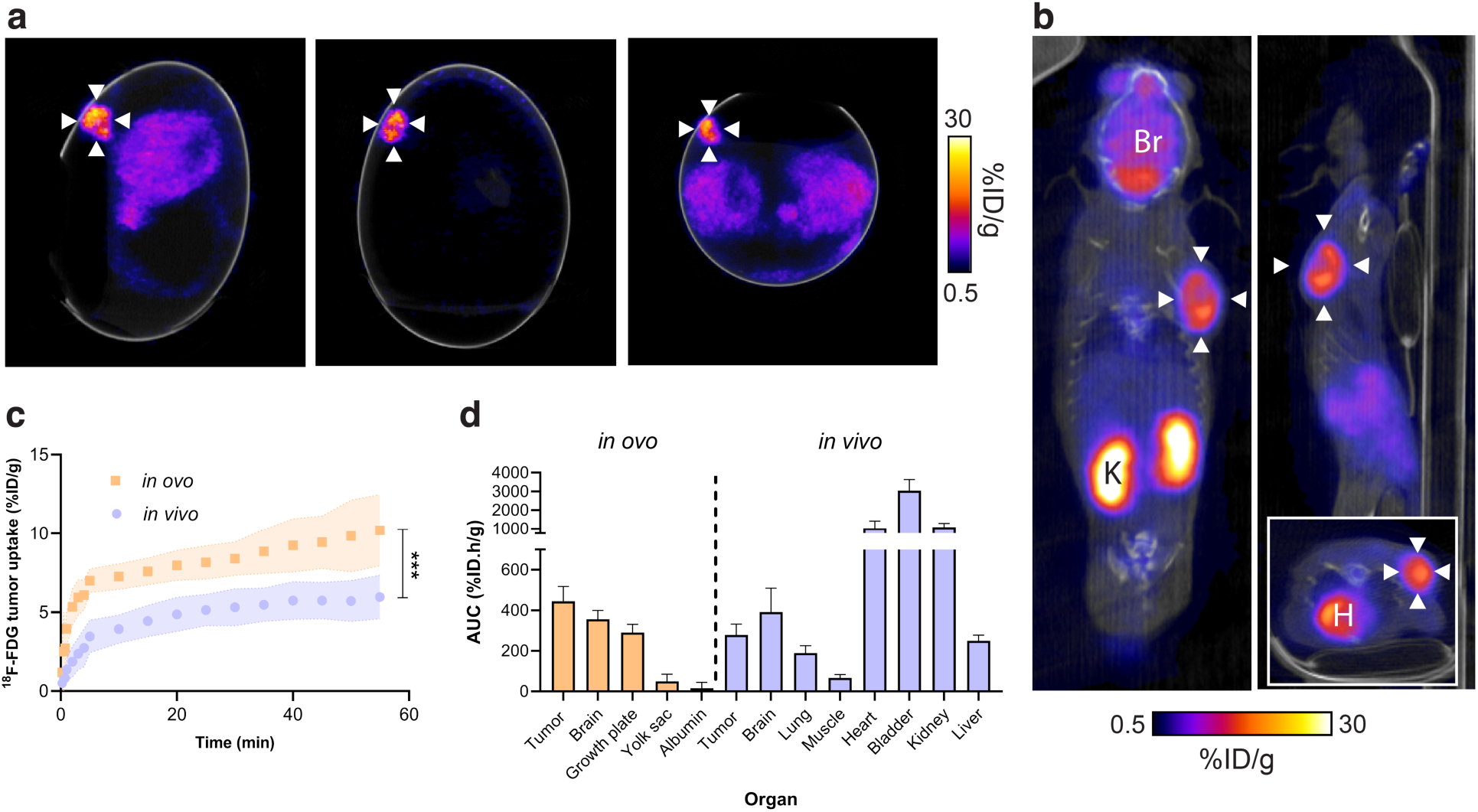
Comparison of *in ovo* and *in vivo* ^18^F-FDG PET/CT imaging. a. Representative *in ovo* ^18^F-FDG PET/CT images 40-60 min p.i.. White arrows indicate the tumor. b. Representative *in vivo* sagittal, coronal and axial (insert) ^18^F-FDG PET/CT images 40-60 min p.i.. White arrows indicate the tumor. Br, brain; H, heart; K, kidney. c. Comparison of *in ovo* and *in vivo* ^18^F-FDG tumor pharmacokinetics. d. *In ovo* and *in vivo* healthy and tumor tissue ^18^F-FDG uptake, expressed as the area under the TAC. Data is expressed as the mean plus standard deviation. *n* = 7 eggs, *n* = 9 mice. ***, *p* < 0.001.

For comparison, dynamic *in vivo* PET/CT imaging was performed in immunocompromised mice implanted with subcutaneous NCI-H460 Fluc xenografts (**Fig. 3b**). Similarly to chick CAM, rapid ^18^F-FDG tumor accumulation occurred over the initial 5 min, followed by a slower rate of retention over the proceeding 55 min (**Fig. 3c**). Tumor-associated radioactivity was lower in the mouse compared to the egg at 60 min p.i., reaching 6.0 ± 1.4 %ID/g; a pattern seen across the entire time course (AUC of 445 ± 73.5 %ID.h/g and 278 ± 53.1 % ID.h/g for *in ovo* and *in vivo* tumors, respectively; *p* = 0.0001; *n* = 9 mice and 7 eggs; **Fig. 3d**). Normal ^18^F-FDG healthy tissue distribution was observed in these mice, characterized by renal excretion, accompanied by high retention in the heart and brain (**Fig. 3d** and **Supplementary Fig. 3c**).

### *In ovo* NCI-H460 Fluc tumors have high but variable ^18^F-FSPG retention

Given the high-quality PET images obtained from the chick CAM, we next assessed ^18^F-FSPG *in ovo* (**Fig. 4a**); a radiotracer with favorable imaging properties whose retention is sensitive to redox manipulations [27]. Tumors exhibited high but variable retention of ^18^F-FSPG (12.7 ± 5.8 %ID/g at 60 min p.i) with a tumor to background ratio of 74 (**Fig. 4b**). ^18^F-FSPG retention was highest in the kidneys, with liver uptake comparable to the tumor (**Fig. 4c**), as confirmed by *ex vivo* biodistribution (**Fig. 5**). Protein expression of the light-chain subunit of system xc^-^, xCT, in the tumor was variable despite minimal changes in the redox-sensitive transcription factor nuclear factor E2-related factor 2 (NRF2; **Fig. 4d**). ^18^F-FSPG tumor retention, however, did not corrolate to GSH (R^2^ = 0.02; *p* = 0.73) or tumor weight (R^2^ = 0.34, *p* = 0.06; **Supplementary Fig. 4**).

**FIGURE 4.**
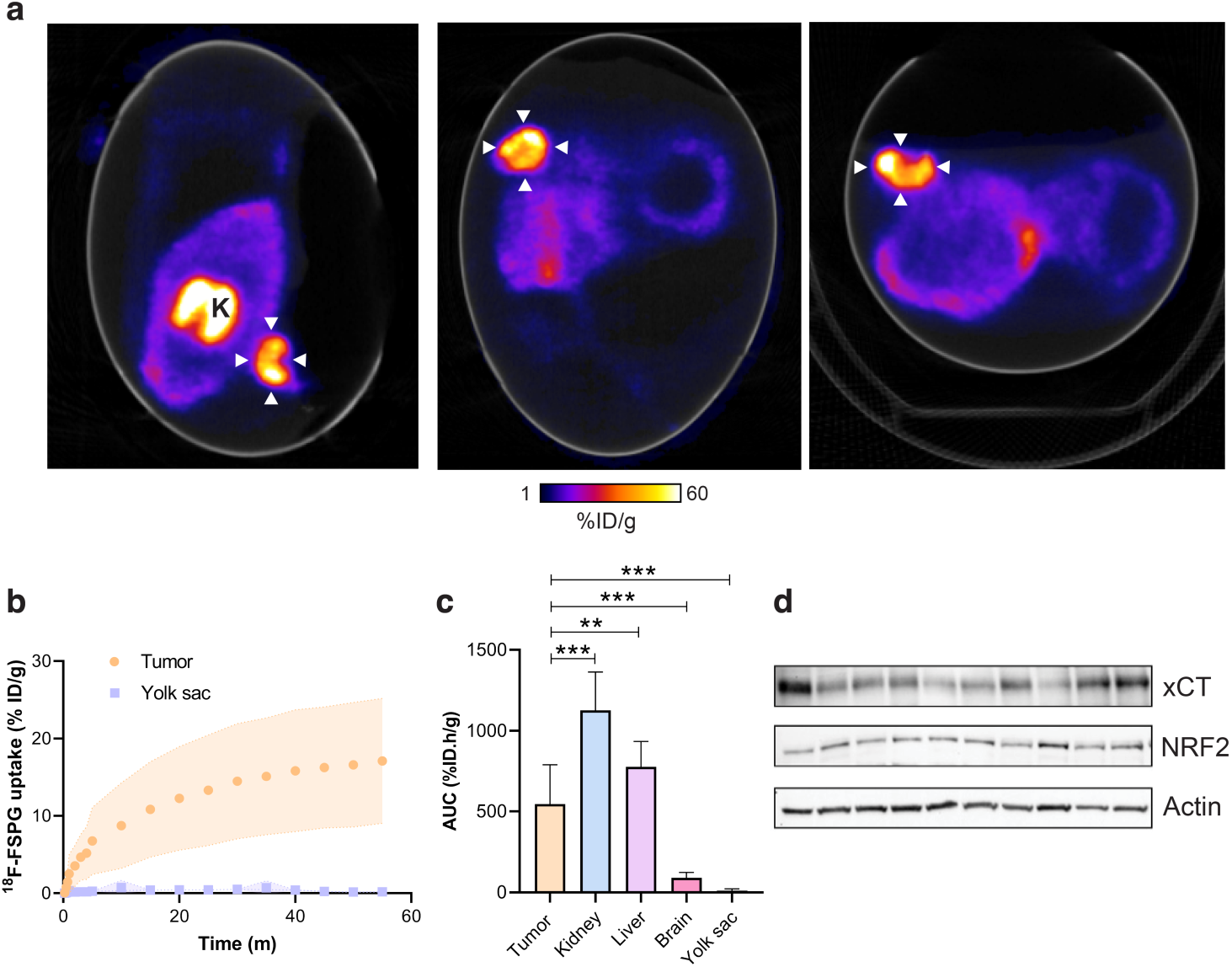
Dynamic ^18^F-FSPG PET imaging *in ovo*. a. Representative *in ovo* ^18^F-FSPG PET/CT images 40-60 min p.i.. White arrows indicate the tumor. K, kidney. b. TAC for tumor and yolk sac-associated ^18^F-FSPG retention *in ovo*. c. AUC for major organs. Data is expressed as a mean plus standard deviation. *n* = 10. d. xCT and NRF2 protein expression from NCI-H460 Fluc *in ovo* tumors. **, *p* < 0.01; ***, *p* < 0.001.

**FIGURE 5.**
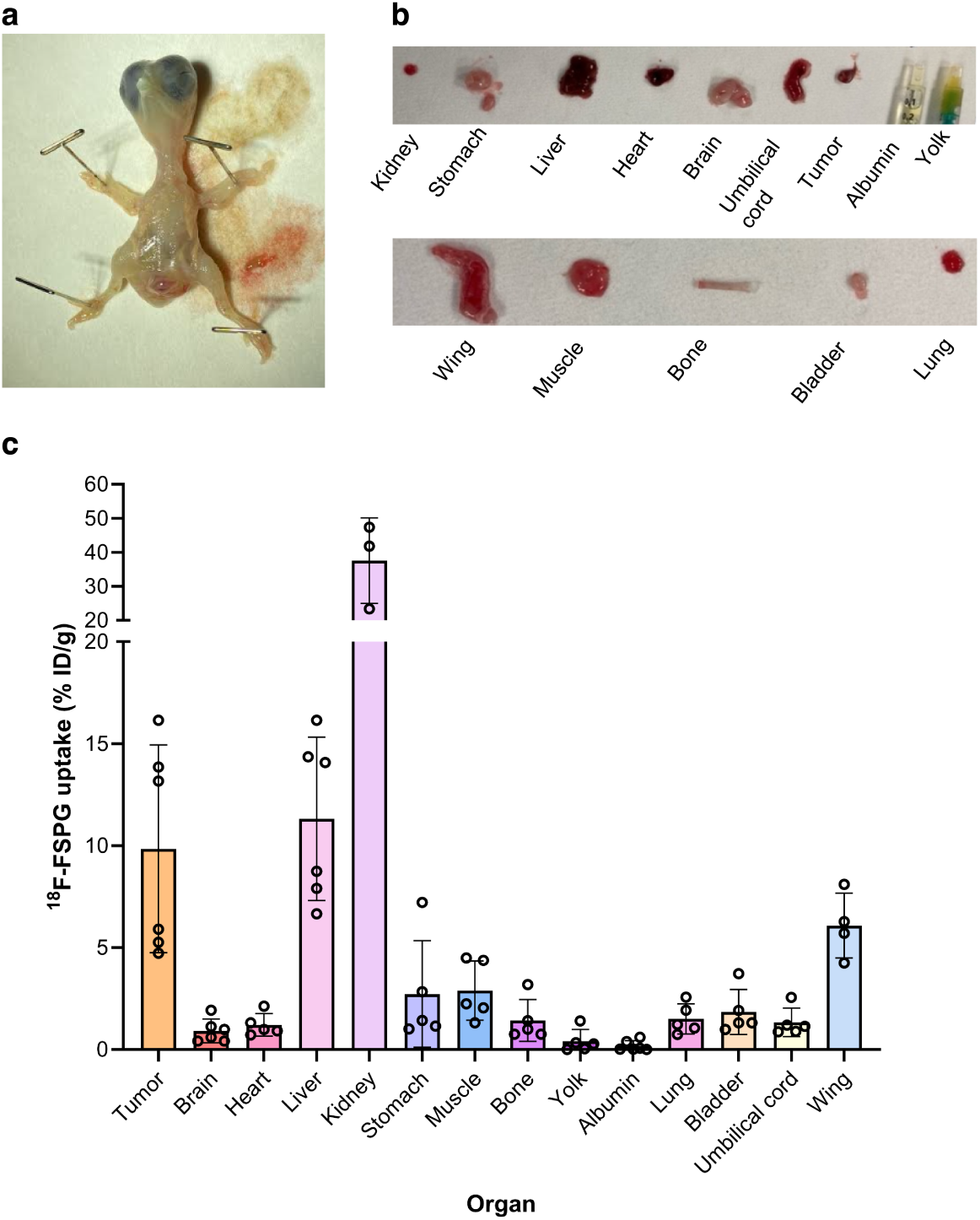
*Ex vivo* chicken embryo biodistribution with ^18^F-FSPG. a. Photo of the excised chick embryo. b. Photos of key organs from the chick embryo at E14. c. ^18^F-FSPG retention in key organs and in NCI-H460 FLuc tumors 60 min p.i. *n* = 3-6.

**FIGURE 6.**
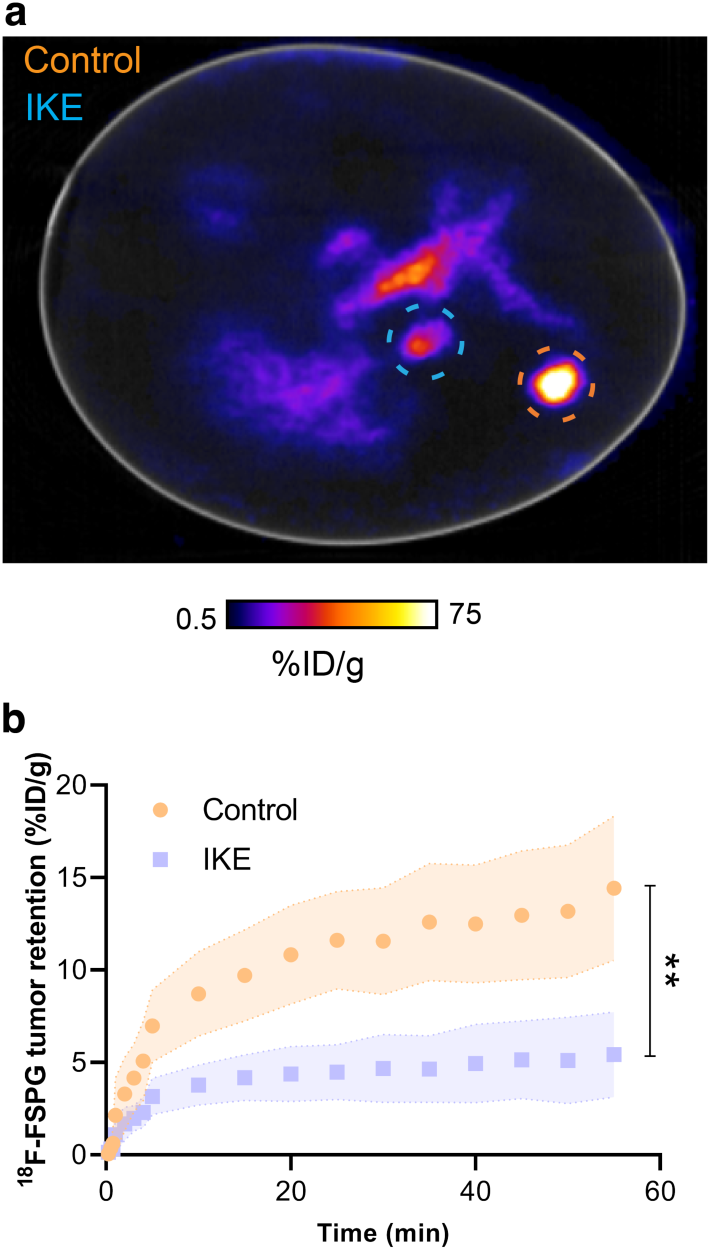
Inhibition of system xc^-^ reduces ^18^F-FSPG uptake. a. Representative ^18^F-FSPG PET/CT image of an egg bearing both control and IKE-treated NCI-H460 Fluc *in ovo* tumors 40-60 min p.i.. Orange circle shows the location of control tumor; blue circle shows the location of IKE-treated tumor. b. ^18^F-FSPG TAC of control *vs*. IKE-treated tumors. Error bars represent one STD from the mean value. *n* = 6; **, *p* = 0.004.

### ^18^F-FSPG uptake is reduced by system xc^-^ inhibition

To assess the utility of the chick CAM in mechanistic imaging studies, we treated eggs bearing NCI-H460 Fluc tumors with an intratumoral injection of the system xc^-^ inhibitor IKE 60 min prior to imaging with ^18^F-FSPG PET/CT (**Fig. 6a**). As we showed previously, high ^18^F-FSPG retention was present in the control tumors (14.4 ± 3.9 %ID/g at 60 min p.i) which was reduced by ∼60% in IKE-treated tumors (5.5 ± 2.3 %ID/g; *n* = 6; *p* = 0.004; **Fig. 6b**).

### ^18^F-FSPG uptake is reduced 24 h after external beam radiotherapy

We, and others, have previously shown that ^18^F-FSPG is an early and sensitive marker of chemotherapy treatment response [28]. To better-understand whether the chick CAM can be used for such applications, we treated NCI-H460 Fluc tumors with 12 Gy external beam x-ray radiotherapy. 24 h after treatment, ^18^F-FSPG retention in treated tumors was reduced by 40 % compared to controls (15.3 ± 6.5 %ID/g and 25.5 ± 7.3 %ID/g, respectively; *n* = 7; *p* = 0.017; **Fig. 7a** & **b**). This decrease coincided with increased tumor apoptosis in treated tumors in the absence of any changes in GSH (**Fig. 7c & d**).

**FIGURE 7.**
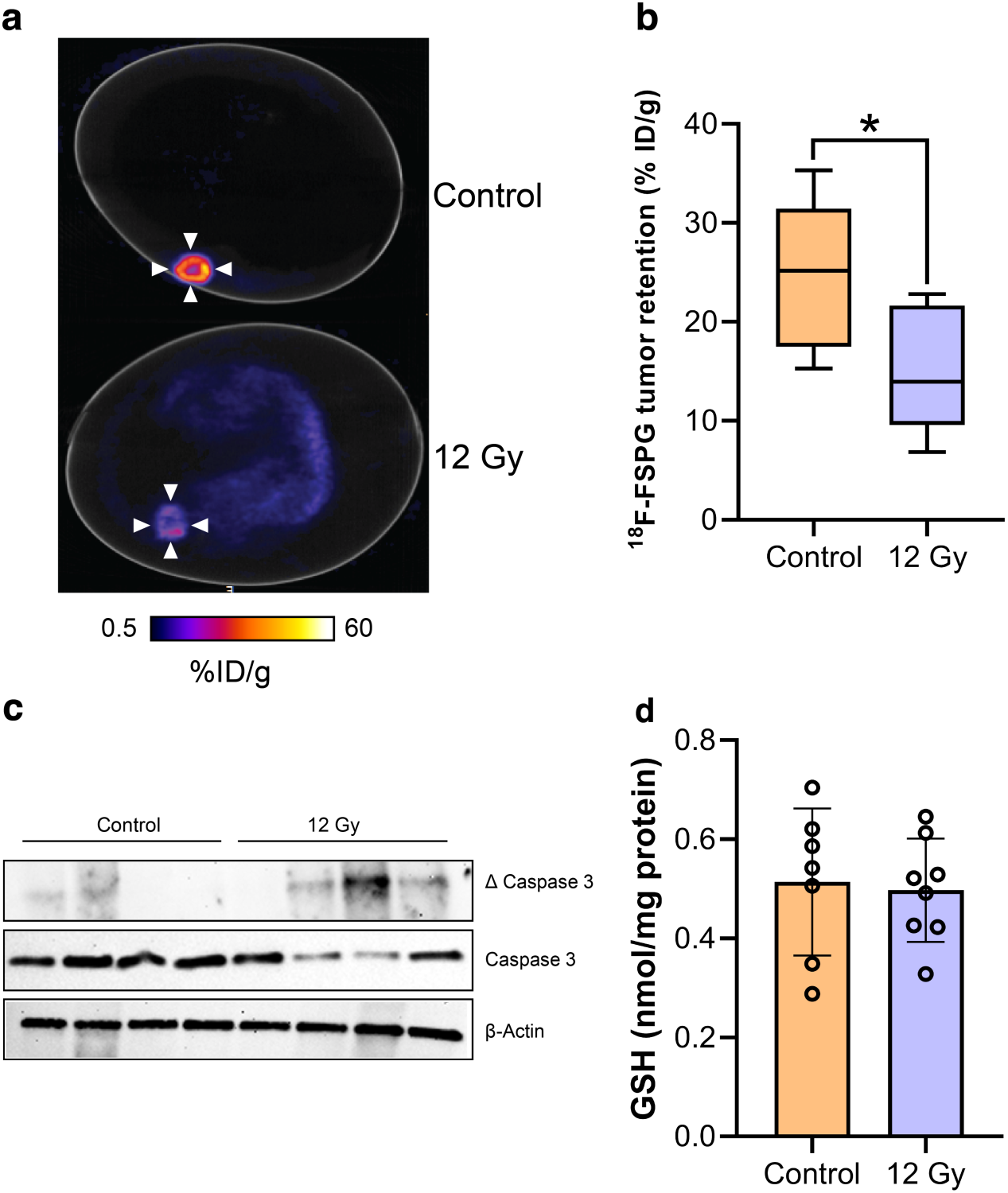
External beam radiotherapy decreases ^18^F-FSPG retention. a. Representative 40-60 min ^18^F-FSPG PET/CT image of NCI-H460 Fluc-bearing eggs treated with 12 Gy radiotherapy or CT alone. b. Quantification of ^18^F-FSPG tumor retention 40-60 min p.i.. *n* = 7; *, *p* = 0.017. c. Western blot showing viability of control and radiation-treated NCI-H460 Fluc *in ovo* tumors. Total and cleaved (Δ) caspase 3 were assessed, with actin used as a loading control (*n* = 4). d. GSH concentrations for control and radiation-treated tumors. *n* = 8.

## DISCUSSION

Animal models of cancer have revolutionized our understanding of cancer. Our ability to recreate this disease using mouse models has contributed to the clinical development of drugs and diagnostic imaging agents [29]. Along the imaging agent development pipeline, however, there are high rates of attrition, leading to significant research and development costs. PET imaging with the chick CAM could potentially minimize animal purchase and maintenance charges, increase experimental throughput, and remove the requirement for an animal license. In our case, delivery and care of 10 BALB/c nu/nu mice for one month alone cost approximately £1,400, while the purchase and delivery of 12 fertilized eggs was £45 with zero maintenance costs: a 97% saving. We found the chick CAM simple to handle and quick to set up, making it an attractive model for those without prior animal training experience. Whilst mammalian models are required to assess radiotracer phamacokinetics, the chick CAM may have utility as an early imaging agent screening tool prior to assessment in more complex disease models.

NSCLC cell lines have previously been shown to grow well *in ovo* [30]. Here, we optimized the growth of the NSCLC cell line NCI-H460 Fluc to consistently cultivate tumors large enough for PET imaging studies. Although ring supports have been used successfully in previous *in ovo* PET imaging studies [31], the plastic rings used here were too heavy for the CAM and introduced infection. Lighter rings made from silicone may be more suitable for use in future studies [31]. Applying trypsin directly onto the CAM (and corresponding enzymatic degration) allowed the tumour cells to embed into the membrane; however, this technique increased cell dispersal and ultimately the development of smaller tumours. Cell suspensions mixed with Matrigel resulted in high tumor take-rates and consistent tumor sizes whilst maximizing embryo viability, as been shown previously [32]. Matrigel contains growth factors (e.g. fibroblast growth factor) which aids proliferation through the promotion of angiogenesis and functions as a solid support for cell engraftment on to the CAM in a suitable location for imaging. Tumors were well-perfused, intersperced with heterogenous areas of necrosis, mimicking what is seen clinically.

Having established a reproducible method for tumor growth with Matrigel, our next aim was to develop a simple cannulation technique to facilitate dynamic *in ovo* PET imaging. The use of glass needles removed the need to perform microsurgery, a method which has hampered previous studies and prevented its wider adoption [22]. The CAM vessels are of an equivalent size to a mouse tail vein, but as these vessels are unsupported (the CAM sits above fluid structures such as the amniotic fluid and yolk sac), an ultra-sharp needle was therefore required.

To determine whether the chick CAM is a viable alternative to simple murine models for preclinical imaging applications, we compared the uptake and retention of the gold standard clinical radiotracer ^18^F-FDG *in ovo* and *in vivo*. We found a similar ^18^F-FDG uptake profile in the chick embryo compared to previous studies [33]. Due to the growth demands at this stage of development, glucose metabolism in many tissues is increased, and as a consequnce we saw high uptake thoughout the embryo, rather than in discrete organs as is typically seen in both mice and humans. Excellent tumor-to-background contrast was achieved in the chick CAM, which followed the same pattern of uptake in mice. ^18^F-FDG tumor uptake *in ovo*, however, was consistently higher than *in vivo*. This variation could be explained by slower ^18^F-FDG blood clearence in the chick embryo compared to the mouse. ^18^F-FDG is rapidly renally excreted *in vivo*, while *in ovo* there is no external clearence pathway [8], which may increase CAM blood radioactivity concentrations and therefore tumor delivery [22]. Our findings are in contrast to recently-published data, where the tumor pharmacokinetics of ^18^F-siPSMA-14 differed between eggs and mice [22]. We hypothesize that these differences are due to the immobilization protocol. Here, we used liquid narcotic medetomidine as opposed to cooling, which can lower metabolic rate and other processes governing radiotracer uptake [31, 34].

Having established a reproducible protocol for dynamic PET imaging in the chick CAM, we next assessed its performance using a variety of different applications with the redox imaging agent, ^18^F-FSPG [35, 36]. High yet variable ^18^F-FSPG retention was measured in the CAM-grown tumors (**Fig. 4**). This variability was not correlated with tumor size, nor was it associated with altered levels of GSH, a surrogate marker of redox status. xCT protein expression varied across chick CAM tumors, providing a possible explanation for the spread of the imaging data. We next used the chick CAM to evaluate target engagement, with pharmacological inhibition of xCT leading to a 60% decrease in ^18^F-FSPG uptake; a decrease comparable to similar experiments performed in mice [27]. These findings highlights the opportunity to use the chick CAM as a high-throughput model for compound screening, and when combined with an appropriate companion diagnostic, mechanistic insight. Changes in uptake of ^18^F-FSPG has been shown to be early indicator of treatment response [27, 28, 36, 37]. Here, 12 Gy of external beam radiation decreased ^18^F-FSPG *in ovo* tumor retention by ∼45% just 24 h after treatment, coinciding with an increase in tumor apoptosis. Chick viability was not impacted by radiotherapy, making the chick CAM an attractive option for treatment response assessment with current and emerging radiotracers.

While this model has several important advantages, no model is without limitations. Firstly, a lack of commercially-available antibodies severely restricts the biochemical analyses that can be performed *ex ovo*. Here, we circumvented this issue by using human tumor xenografts and an *in ovo* method of vascular staining which doesn’t require antibody labeling. Additionally, at E14 most of the structures in the egg are soft and CT cannot be used to draw ROIs. Consequently, ROIs are drawn on the PET signal, increasing the chances of mischaracterization due to spill-over effects from the vasculature. To overcome this issue, CT contrast agents may help delineate tissue boundaries [14], with anatomical imaging dramatically improved with PET-MRI [22]. In addition to issues related to vessel cannulation described above, egg-to-egg differences in vessel formation means that some eggs are not suitable for cannulation, leading to wastage. Unfortunately, we found that removal of injection cannulae often results in heavy bleeding, complicating serial intravenous radiotracer injections which might otherwise be used for longitudinal imaging [38]. Repeat drug administration, however, is possible through delivery *via* the yolk sack [39]. In our study, intra-tumoral injections were selected to minimize drug metabolism and reduce toxicity to the chick, but there remains a flexibility in possible approaches. Finally, development of a multi-egg bed for the PET scanner, similar to the mouse hotel [4], would additionally increase throughput and further decrease costs.

## CONCLUSION

We have shown it possible to reproducibly cultivate *in ovo* NSCLC tumors for imaging just seven days after implantation. Dynamic PET imaging of these tumors was possible using a simple cannulation method without the requirement for microsurgery. Chick CAM tumors were avid for both ^18^F-FDG and ^18^F-FSPG, providing high signal-to-background ratios. This work supports the case for the use of the chick CAM as a more sustainable, low-cost substitute to tumor xenograft mouse models and has the potential to both expedite novel radiotracer development and assess tumor response to treatment.

## Abbreviations

(*S*)-4-(3-^18^F-fluoropropyl)-ʟ-glutamate, ^18^F-FSPG; ^18^F-2-fluoro-2-deoxy-ᴅ-glucose, ^18^F-FDG;; Analysis of variance, ANOVA; glutathione, GSH; maximum-intensity projections, MIPs; positron emission tomography, PET; post injection, p.i.; standard deviation, SD. Imidazole-Ketone erastin, IKE.

## Acknowledgements

The authors would like to thank Professor Michael Shattock for advice on cannulation and donation of multiple pieces of equipment, Dr Jana Kim and Dr Kavitha Sunassee for practical help with *in vivo* experiments, and Mr David Thakor and Dr Matt Hutchings for setting up the egg facility in the School of Biomedical Engineering & Imaging Sciences at King’s. For the purpose of open access, the authors have applied a CC BY public copyright license to any Author Accepted Manuscript version arising from this submission.

