## Supplemental data for "The Chicken Chorioallantoic Membrane as a Low-Cost, High-Throughput Model for Cancer Imaging"

### SUPPLEMENTAL FIGURES AND LEGENDS

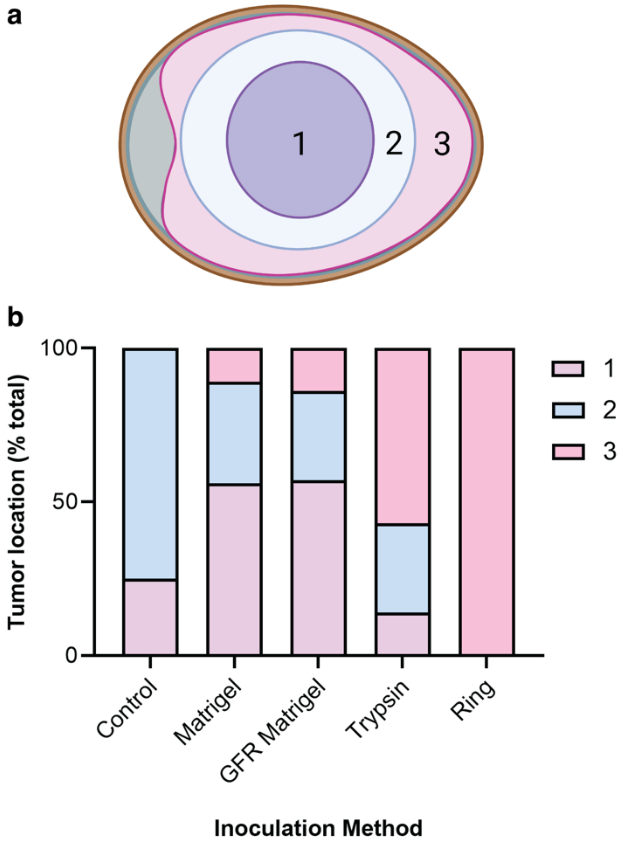


**SUPPLEMENTAL FIGURE 1.** Location of chick CAM tumor growth with various inoculation methods. a. Schematic illustrating the possible positions of tumor growth within the egg. Location 1: the center of the CAM. Location 2: the edges of the CAM. Location 3: the inside of the shell. b. Location of grown tumors based on inoculation method and matrix.

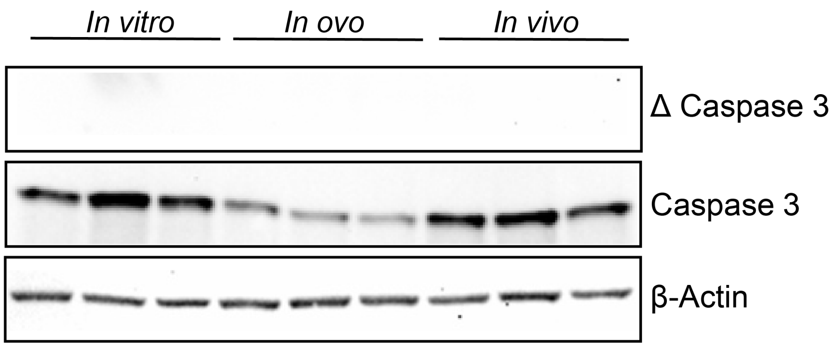


**SUPPLEMENTAL FIGURE 2.** Tumors grown *in ovo* have low baseline levels of apoptosis. Cleaved caspase 3 levels (Δ) were assessed in NCI-H460 Fluc cells grown *in vitro*, *in ovo*, and *in vivo*. Actin and total caspase 3 were used as controls.

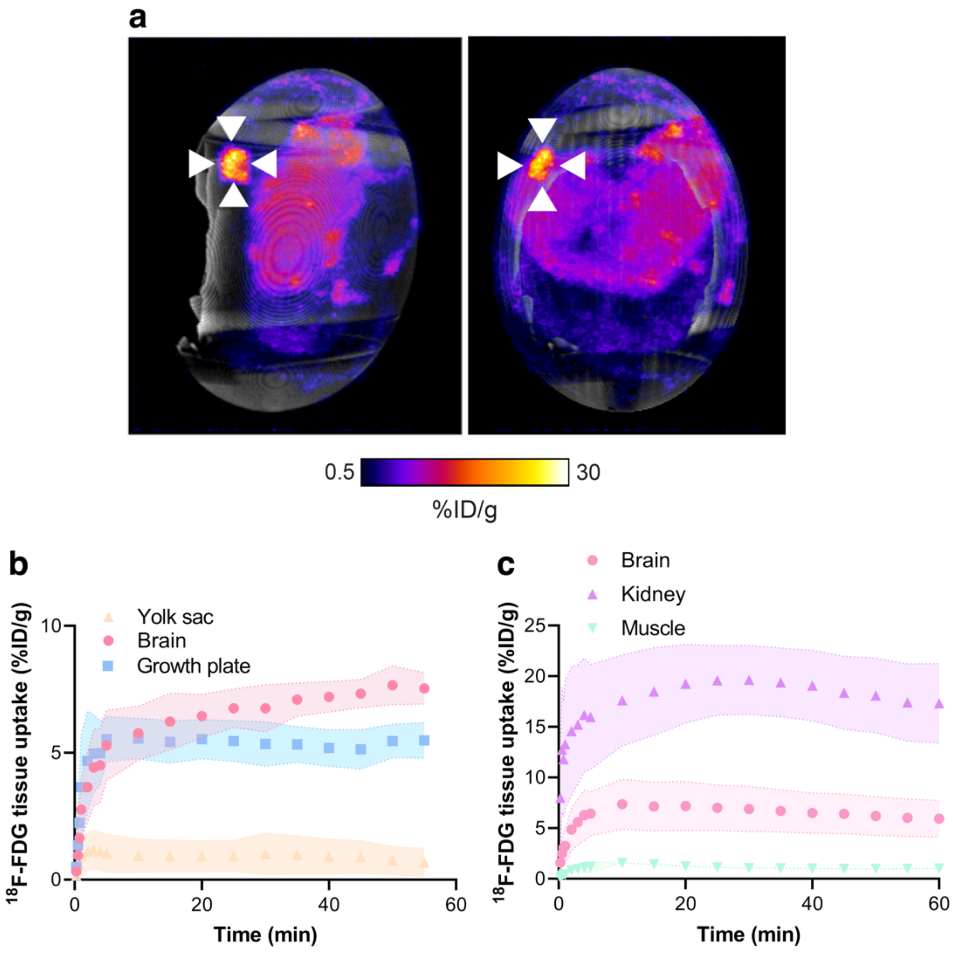


**SUPPLEMENTAL FIGURE 3.** $^{18}$F-FDG PET/CT maximum intensity projections and healthy tissue time activity curves. a. $^{18}$F-FDG PET/CT maximum intensity projection (0 – 60 min p.i.) from a chick CAM with an NCI-H460 Fluc xenograft tumor (white arrow heads). b. TAC for $^{18}$F-FDG tissue uptake from the organs of the embryo. *n* = 7. c. TAC for $^{18}$F-FDG tissue uptake in mice. *n* = 9. Shaded regions represent one standard deviation.

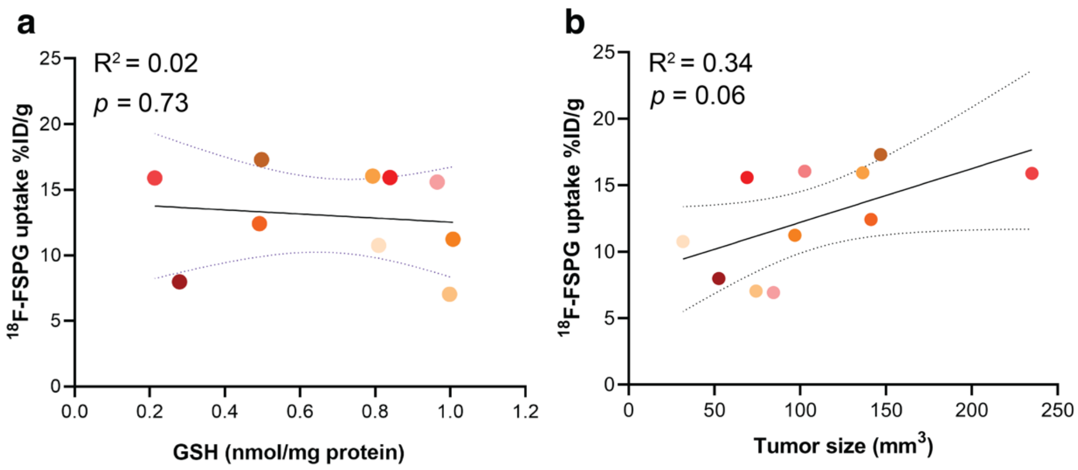


**SUPPLEMENTAL FIGURE 4.** $^{18}$F-FSPG retention in the chick CAM doesn't correlate with tumor size or GSH concentration. a. Correlation plot of tumor $^{18}$F-FSPG retention *vs.* intracellular GSH. b. Correlation plot of tumor size *vs.* tumor $^{18}$F-FSPG retention.
